# Primate TRIM34 is a broadly-acting, TRIM5-dependent lentiviral restriction factor

**DOI:** 10.1101/2023.03.24.534139

**Authors:** Joy Twentyman, Anthony Khalifeh, Abby L. Felton, Michael Emerman, Molly OhAinle

## Abstract

Human immunodeficiency virus (HIV) and other lentiviruses adapt to new hosts by evolving to evade host-specific innate immune proteins that differ in sequence and often viral recognition between host species. Understanding how these host antiviral proteins, called restriction factors, constrain lentivirus replication and transmission is key to understanding the emergence of pandemic viruses like HIV-1. Human TRIM34, a paralogue of the well-characterized lentiviral restriction factor TRIM5α, was previously identified by our lab via CRISPR-Cas9 screening as a restriction factor of certain HIV and SIV capsids. Here, we show that diverse primate TRIM34 orthologues from non-human primates can restrict a range of Simian Immunodeficiency Virus (SIV) capsids including SIV_AGM-SAB_, SIV_AGM-TAN_ and SIV_MAC_ capsids, which infect sabaeus monkeys, tantalus monkeys, and rhesus macaques, respectively. All primate TRIM34 orthologues tested, regardless of species of origin, were able to restrict this same subset of viral capsids. However, in all cases, this restriction also required the presence of TRIM5α. We demonstrate that TRIM5α is necessary, but not sufficient, for restriction of these capsids, and that human TRIM5α functionally interacts with TRIM34 from different species. Finally, we find that both the TRIM5α SPRY v1 loop and the TRIM34 SPRY domain are essential for TRIM34-mediated restriction. These data support a model in which TRIM34 is a broadly-conserved primate lentiviral restriction factor that acts in tandem with TRIM5α, such that together, these proteins can restrict capsids that neither can restrict alone.

## INTRODUCTION

Restriction factors are a class of cell-intrinsic, germline-encoded host immune factors that can inhibit viral infection and replication. The human genome encodes for approximately 70-100 TRIM (Tripartite Motif) proteins, many of which play a role in host defense [1]. The alpha isoform of TRIM5 (TRIM5α) in primates is a well-characterized example of a primate restriction factor with activity against retroviruses. The activity of TRIM5α against a given retrovirus depends on both the species from which the TRIM5α is derived and the capsid (CA) protein of the retrovirus [2–5]. In the context of HIV-1, TRIM5α acts directly on CA by multimerizing onto the CA lattice [5,6]. This interaction results in aberrant uncoating of CA, interrupting the viral life cycle [5,7]. TRIM5α from rhesus macaques and many other Old World monkeys has much greater antiviral activity against HIV-1 than does human TRIM5α [2,5,8].

TRIM proteins are composed of a common set of N-terminal domains (RING, Bbox, and coiled-coil) followed by one or more variable C-terminal domains [9]. An important characteristic of TRIM5α, and TRIM proteins more generally, is their ability to oligomerize to form structures with very high binding avidity to CA [7]. Homo-oligomerization of TRIM5α is essential to its ability to restrict viral CA [10–12]. Furthermore, TRIM proteins have also been demonstrated to hetero-oligomerize with each other [9,13,14].

Previously, our lab performed a CRISPR-Cas9 screen to identify restriction factors against an HIV-1 strain with a mutation in CA, N74D, that renders it more susceptible to CA-mediated restriction factors [15]. We identified TRIM34 as a restriction factor of the HIV-1 N74D capsid mutant as well as select SIV capsids [16]. TRIM34 is a paralogue of TRIM5α, sharing a common domain architecture, and human TRIM34 and TRIM5α have approximately 57% amino acid identity [17]. Moreover, TRIM34-mediated restriction requires TRIM5α, and TRIM34 and TRIM5α colocalize with incoming capsids [16].

While TRIM5α and TRIM34 both can interact with lentiviral CA, TRIM5α, but not TRIM34, has undergone positive selection [14,18]. A history of positive selection, in which the rate of nonsynonymous mutations exceeds the rate of synonymous mutations, is characteristic of host proteins that are in evolutionary conflict with pathogens and often occurs at sites of direct physical interface between host and pathogen [19,20]. Specifically, the v1 loop of TRIM5α’s C-terminal SPRY domain has been found to have undergone rapid evolution, and this region is responsible for viral recognition [18]. Conversely, the lack of evidence for positive selection on TRIM34 suggests that TRIM34 may not contain sites of evolutionarily-important direct viral interaction.

Although currently there does not exist evidence of evolutionary conflict within TRIM34, here we show that TRIM34 antiviral activity has been broadly conserved across primate species as an antiviral gene against CA of diverse primate lentiviruses. We find that restriction by TRIM34 from primates requires TRIM5α and that the TRIM5α SPRY v1 loop is an essential mediator of restriction. These results suggest that TRIM34 relies on the capsid-binding properties of TRIM5α. However, as the TRIM34 SPRY domain is also required for restriction, our results suggest that both TRIM34 and TRIM5α contribute to capsid recognition and/or antiviral function. We propose that TRIM34, which has not undergone positive selection, is an antiviral protein that requires TRIM5α, which has undergone positive selection.

## RESULTS

### Diverse primate TRIM34 orthologues can act as lentiviral restriction factors

Amino acid variation due to positive selection over evolutionary time in the v1 loop of TRIM5α confers different primate species with varying specificities towards retroviral capsids [18,21]. Although TRIM34 lacks evidence of positive selection found in TRIM5α, there is still diversity across different primate TRIM34 alleles. For example, relative to human TRIM34, chimpanzee, sabaeus monkey, and rhesus macaque orthologues differ by 4, 27, and 26 amino acids, respectively. While we have previously shown that human TRIM34 restricts SIV_AGM-TAN_ and SIV_MAC_, here we wished to determine whether amino acid variation in TRIM34 orthologues from divergent primates affects TRIM34’s capacity to restrict different primate lentiviral capsids [16]. Specifically, we tested whether a panel of 4 TRIM34 orthologues from diverse primate species comprising human (*Homo sapiens*), chimpanzee (*Pan troglodytes troglodytes*), sabaeus monkey (*Chlorocebus sabaeus*), and rhesus macaque (*Macaca mulatta*) orthologues would restrict a panel of lentiviral capsids (HIV-1, SIV_CPZ_, SIV_AGM-SAB_, SIV_AGM-TAN_, and SIV_MAC_, which arise from humans; chimpanzees; two species of African green monkeys, the sabaeus monkey and the tantalus monkey; and rhesus macaques, respectively).

To test the effects of different primate TRIM34 orthologues in the absence of endogenous TRIM34, we generated a clonal *TRIM34* knockout (KO) line in the human THP-1 cell line. We then introduced primate TRIM34 orthologues using a doxycycline-inducible lentiviral vector to generate human, chimpanzee, sabaeus, and macaque TRIM34 over-expressing cell lines. These orthologues were selected because they represent a broad range of primate TRIM34 from both hominid and Old World monkeys for which we also have the corresponding lentiviral capsids. We tested an empty-vector line as a control for the absence of any TRIM34. We confirmed that expression of each of the TRIM34 orthologues is induced by doxycycline (Figure 1a), although overall steady-state expression levels varied somewhat with the macaque TRIM34 expressed at higher levels and the chimpanzee TRIM34 at lower levels.

**Figure 1.**
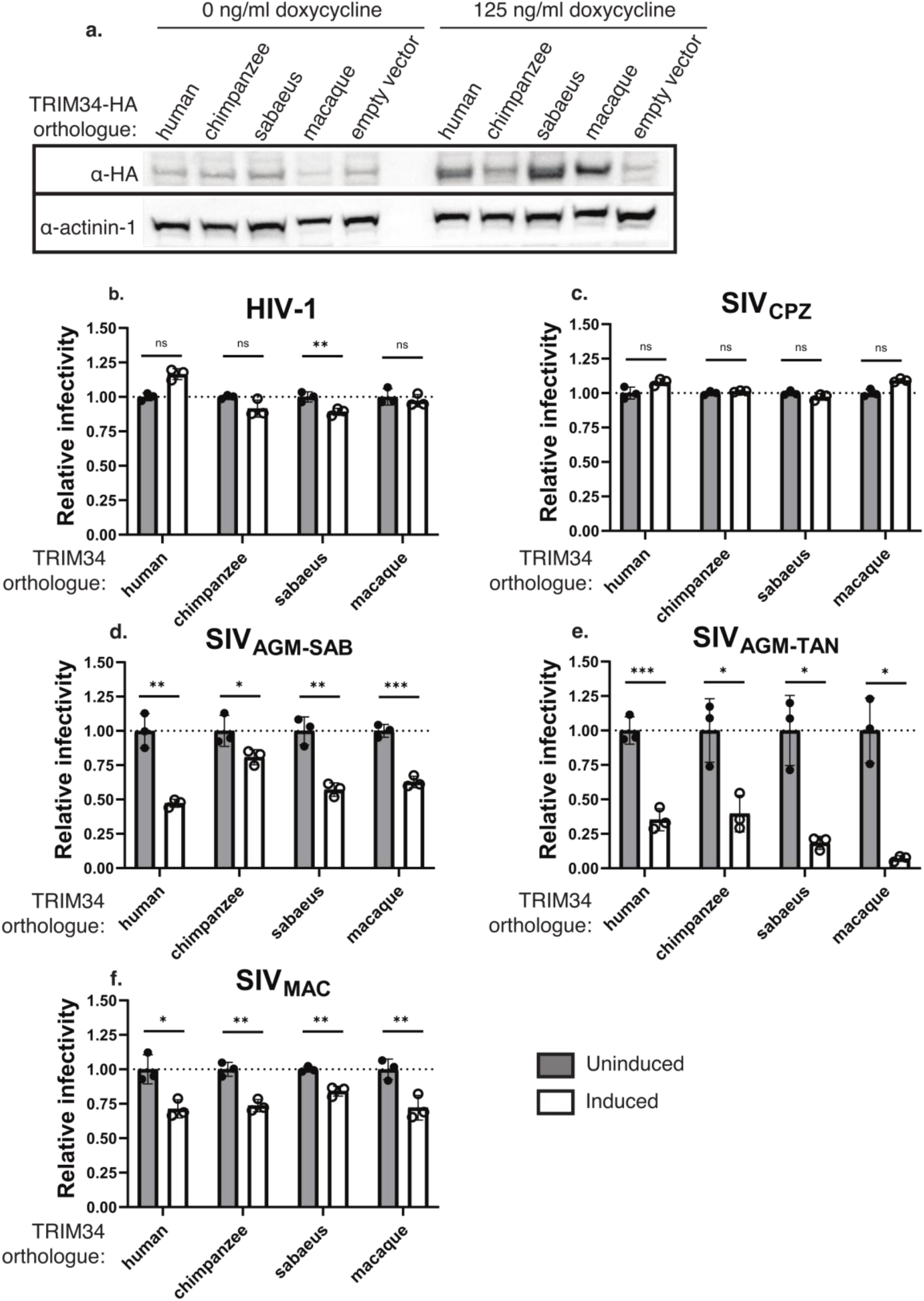
Diverse primate TRIM34 orthologues restrict SIV_AGM-SAB_, SIV_AGM-SAB_, and SIV_MAC_ capsids in the presence of TRIM5α. a. THP-1 *TRIM34* clonal KO cells were generated by electroporation of multiplexed sgRNA against *TRIM34* and single cell sorting (ICE KO score = 96%). THP-1 *TRIM34* KO cells were transduced with doxycycline-inducible HA-tagged primate TRIM34 orthologues or empty vector control. Primate TRIM34 expression was induced with doxycline. Expression levels were visualized by immunoblotting 72 h post-induction. b-f. Primate TRIM34 expression was induced in THP-1 *TRIM34* clonal KO cells. 1 day post-induction, cells were either challenged with chimeric virus particles containing HIV-1 capsids co-expressing zsGreen (b) or SIV capsids co-expressing eGFP including SIV_CPZ_ (c), SIV_AGM-SAB_ (d), or SIV_MAC_ (f); or challenged with VSV-G pseudotyped SIV_AGM-TAN-luc_ (e). Infection was quantified 2 dpi by flow cytometry (b-d, f) or luminometry (e). Relative infectivity in induced cells (white bars) is normalized to average infectivity in uninduced control cells (grey bars). Data are represented as mean +/- s.d. of 3 technical replicates from 1 representative experiment. One-sided p values were calculated by Welch’s *t*-test. ns = nonsignificant, * p ≤ 0.05, ** p ≤ 0.01, *** p ≤ 0.001, **** p ≤ 0.0001.

We then challenged these cells to infection using a VSV-G pseudotyped lentiviral vector system, in which the CA region of a gag/pol expression construct encoded either HIV-1, SIV_CPZ_, SIV_AGM-SAB_, or SIV_MAC_ capsids, whose host species match the species of origin of the TRIM34 orthologues used [22,23]. These virus particles also encoded a fluorescent reporter. We also tested a full-length infectious molecular clone of SIV_AGM-TAN_ that encodes a luciferase reporter [24]. After induction with doxycycline or media-only control, infectivity was quantified 2 days post-infection (dpi) by luciferase assay (SIV_AGM-TAN_) or flow cytometry (all others). Of note, we found that while none of the TRIM34 orthologues tested restricted HIV-1 or SIV_CPZ_ capsids (Figure 1b-1c), all of these same TRIM34 orthologues restricted SIV_AGM-TAN_, SIV_AGM-SAB_, and SIV_MAC_ capsids (Figure 1d-1f). These data suggest that, in contrast to TRIM5α, the antiviral specificity of TRIM34 does not seem to vary across TRIM34 orthologues. Rather, TRIM34 antiviral activity against the same set of lentiviral capsids is a conserved activity of all primate TRIM34 orthologues we tested. Moreover, the antiviral activity of all primate TRIM34s is specific for the same subset of lentiviral capsids. For example neither of the hominid lentiviruses were affected by any TRIM34 orthologue we tested (Figure 1b-c), while the lentiviral capsids that were restricted by human TRIM34—SIV_AGM-TAN_, SIV_AGM-SAB_, and SIV_MAC_—were also restricted by the other primate TRIM34 orthologues. In sum, these data support the hypothesis that TRIM34 restriction is a conserved activity in primates with shared specificity for certain primate lentiviral capsids.

### TRIM34 requires TRIM5α for restriction

Previously, we found that human TRIM34 requires TRIM5α to restrict HIV-1 N74D capsids [16]. Given that the TRIM34 orthologues tested all show selectivity for the same subset of lentiviral capsids (Figure 1), we reasoned that TRIM5α might be responsible for the capsid specificity of TRIM34-mediated restriction. Notably, for the experiments described in Figure 1 in which all TRIM34 orthologues tested restricted the same subset of capsids, the cells used contained endogenous TRIM5α. Therefore, we next asked whether TRIM5α was more broadly required for TRIM34-mediated restriction of SIV_AGM-SAB_, which was the most potently restricted capsid of the capsids tested in Figure 1. Specifically, we assayed for restriction in TRIM34-overexpressing cells in which we remove endogenous TRIM5α expression. We created pooled knockouts of *TRIM5* in the background of THP-1 *TRIM34* clonal KO cells containing doxycycline-inducible human or macaque TRIM34. Although we observed high KO efficiency scores for the pooled KO lines (Figure 2; legend), these pooled KO cell lines still contained some TRIM5α expression as KO is not complete at a population level. We also generated control cell lines using non-targeting control guides (NTCs); these cells still contained endogenous TRIM5α. We then infected these cells with SIV_AGM-SAB_. Relative to cells missing both TRIM34 and TRIM5α (Figure 2a-b; black symbols), SIV_AGM-SAB_ is restricted only in the presence of both TRIM34 and TRIM5α (Figure 2a-b; red symbols). Notably, this was true for both human and rhesus macaque TRIM34 (Figure 2a – human; Figure 2b – macaque). That is, human TRIM5α can fulfill TRIM34’s requirement for TRIM5α, even for a TRIM34 orthologue from a different primate species. Conversely, SIV_AGM-SAB_ is not restricted either by cells expressing only TRIM34 that are knocked out for TRIM5α (Figure 2a-b; green symbols), nor by cells expressing only endogenous TRIM5α without induction of TRIM34 (Figure 2a-b; blue symbols). Although it does appear that there may be a small amount of restriction in the presence of TRIM34 only (Figure 2a-b; green symbols), we attribute this to the fact that the *TRIM5* KO cells were generated as a KO pool: therefore, at a population level, some cells in this cell pool still have intact TRIM5α expression. Overall, these data suggest that primate TRIM34 orthologues broadly require TRIM5α for restriction of lentiviral capsids.

**Figure 2.**
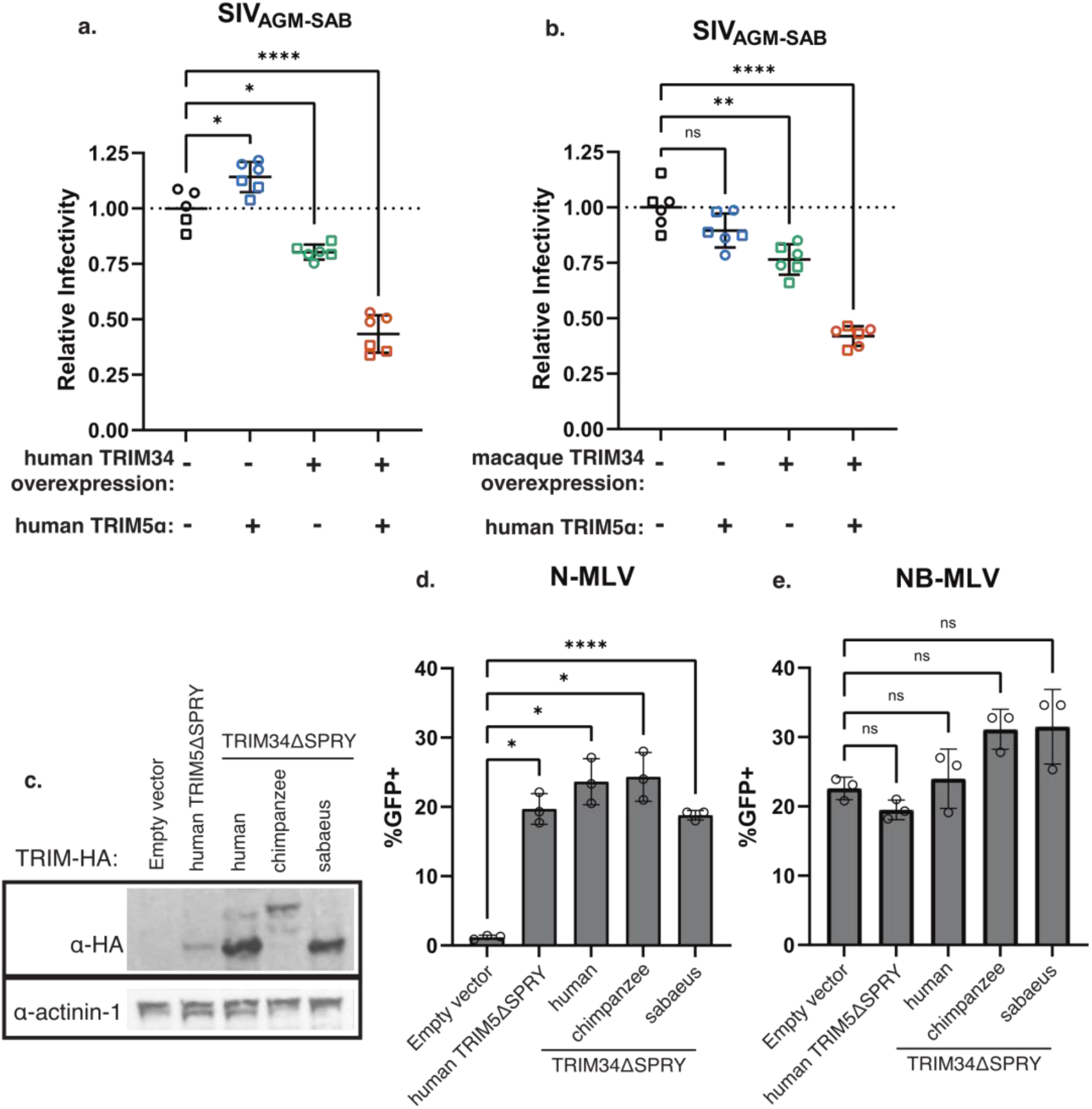
TRIM34 requires TRIM5α to restrict SIV_AGM-SAB_. a-b. THP-1 *TRIM34* clonal KO cells containing doxycycline-inducible TRIM34 from humans (a) or rhesus macaques (b) were transduced with lentiviral vectors encoding Cas9 and 1 of 2 independent sgRNA against *TRIM5* or a non-targeting control to generate pooled *TRIM5* knockout or NTC cell lines. KO efficiency was assessed by ICE analysis. For human TRIM34-expressing cell lines, ICE KO efficiency scores were as follows: *TRIM5* sgRNA 1 (squares) = 76, *TRIM5* sgRNA 2 (circles) = 90. For rhesus macaque TRIM34-expressing cells, ICE KO scores were as follows: *TRIM5* sgRNA 1 (squares) = 83, *TRIM5* sgRNA 2 (circles) = 93. Thus, there existed 4 different cellular conditions: no TRIM34 or TRIM5α expression = black symbols, only endogenous TRIM5α expression = blue symbols, only overexpressed TRIM34 = green symbols, both endogenous TRIM5α and overexpressed TRIM34 = red symbols. 1 day after doxycycline induction, cells were infected with SIV_AGM-SAB_ CA particles; each infection was performed in triplicate. 2 dpi, infectivity was quantified by flow cytometry. Relative infectivity is normalized to mean infectivity in *TRIM5* KO, TRIM34-uninduced cells (black symbols). Data are represented as mean +/- s.d. from 1 representative experiment. Mean represents average across all 6 replicates. c. HeLa cells were transduced to stably express primate TRIMΔSPRY orthologues or empty vector control. Expression was visualized by immunoblotting. d-e. HeLa cells stably expressing primate TRIM34ΔSPRY orthologues or empty vector control were infected with N-MLV (d) or NB-MLV (e). Level of infection was quantified 2 dpi by flow cytometry. Data are represented as mean +/- s.d. from 1 representative experiment. Infections were performed in triplicate. a-b, d-e. p values were calculated by Brown-Forsythe and Welch’s 1-way ANOVA with Dunnett’s T3 test for multiple comparisons. ns = nonsignificant, * p ≤ 0.05, ** p ≤ 0.01, *** p ≤ 0.001, **** p ≤ 0.0001.

### Human TRIM5α functionally interacts with TRIM34 from different primates

We next sought to assess whether TRIM34 orthologues and TRIM5α can functionally interact with each other via a TRIM5α restriction assay. Previous work has shown that dimerization and higher-order multimerization are essential for TRIM5α-mediated restriction [14,25–27]. Furthermore, TRIM5α has been shown to associate not only with itself but also with TRIM34 via two-hybrid screen and co-immunoprecipitation [9,13,14]. To assess functional interaction of TRIM34 and TRIM5α more directly, we exploited the fact that TRIM proteins that have been deleted of their SPRY domains (TRIMΔSPRY) exert a dominant-negative effect on restriction of N-tropic MLV (N-MLV) by endogenous TRIM5α constitutively expressed in HeLa cells [13]. Overexpression of human TRIM34ΔSPRY inhibits restriction of N-MLV in HeLa cells by the endogenous TRIM5α [13]. This implies that TRIM34 is able to functionally interact with TRIM5α. Thus, we asked whether primate TRIM34ΔSPRY orthologues could also interact with human TRIM5α in this assay. The human and rhesus macaque TRIM34ΔSPRY constructs were truncated at the start of the SPRY domain, while the chimpanzee TRIM34ΔSPRY construct was truncated partway through the SPRY domain, between the v2 and the v3 loops. We overexpressed the HA-tagged human, chimpanzee, and macaque TRIM34ΔSPRY constructs in HeLa cells together with human TRIM5ΔSPRY as a positive control (Figure 2c). These cells were then infected with N-MLV. We found that relative to empty vector control, overexpression of all three TRIM34ΔSPRY orthologues, as with TRIM5ΔSPRY, relieved restriction of N-MLV by human TRIM5α (Figure 2d). We also infected with NB-tropic MLV (NB-MLV) as a control (Figure 2e), since NB-MLV is not restricted by TRIM5α [28,29]. These data further support our model that diverse TRIM34 orthologues and human TRIM5α can functionally interact with each other.

### Both TRIM34 and TRIM5α SPRY domains are involved in viral restriction

Since TRIM5α is required for restriction by TRIM34 and does not seem to vary depending on which TRIM34 is used, we hypothesized that capsid recognition comes from TRIM5α, and not from TRIM34. In the case of TRIM5α-mediated restriction, the SPRY domain, and specifically the v1 loop, determines antiviral specificity [18,21,30]. Furthermore, altering a single amino acid in the human TRIM5α SPRY domain (arginine 332) in the v1 loop to match the corresponding macaque residue (proline) is sufficient to confer strong restriction of HIV-1 [21,30]. To ask if the TRIM5α SPRY domain is involved in TRIM34-mediated CA recognition, we used cells that were doubly knocked out for endogenous *TRIM5* and *TRIM34* (THP-1 *TRIM34 TRIM5* clonal KO cells). In these cells, we introduced inducible vectors expressing wild type (WT) human TRIM34 with WT human TRIM5α, TRIM5α in which the v1 loop was deleted (TRIM5α Δv1), and TRIM5α in which the arginine at position 332 was mutated to a proline (TRIM5α R332P). We expressed each TRIM individually as well as in the context of each other and confirmed expression levels by Western blot (Figure 3a). In concordance with the results in Figure 2, we found that SIV_AGM-SAB_ was restricted only in the presence of both human TRIM34 and human TRIM5α, but not when either TRIM34 or TRIM5α were expressed in isolation (Figure 3b; grey bars). Furthermore, in agreement with our previous findings [16], HIV-1 N74D, but not WT HIV-1, was restricted only in the presence of both TRIM34 and TRIM5α (Figure 3c-d; grey bars).

**Figure 3.**
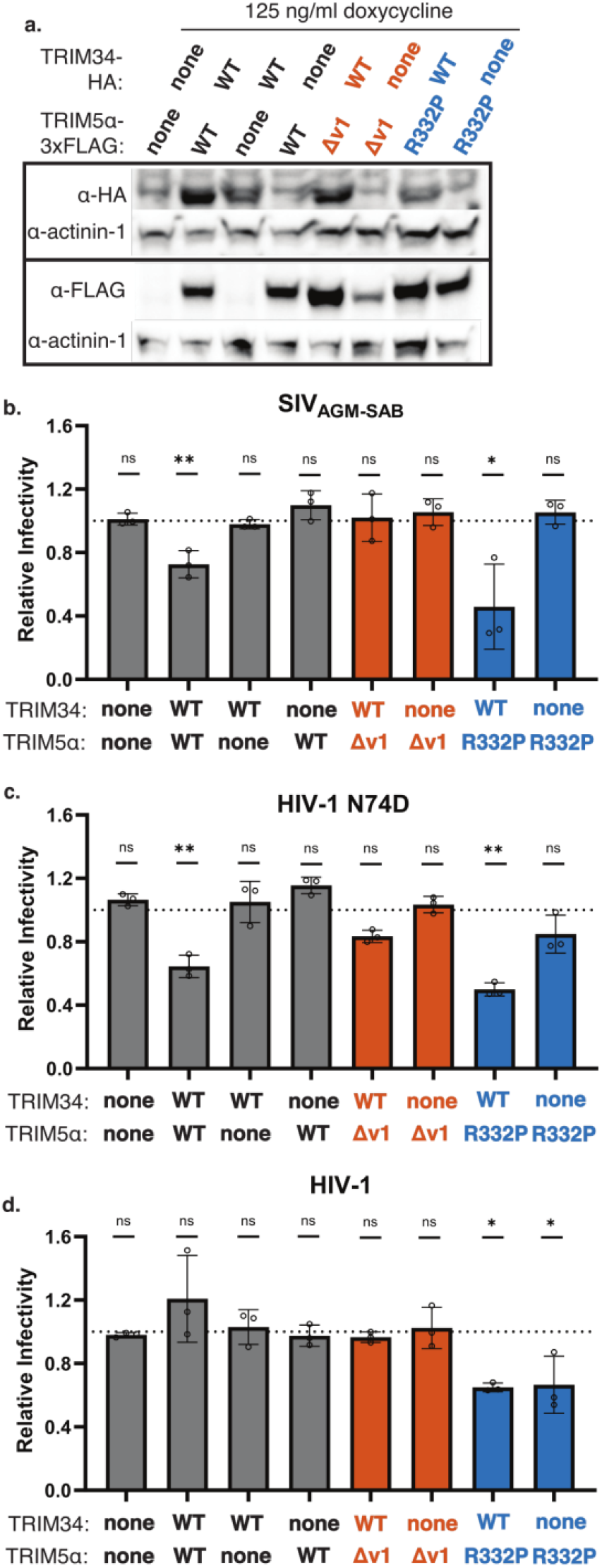
TRIM5α v1 loop is necessary for restriction of SIV_AGM-SAB_. a. THP-1 *TRIM34* clonal KO cells were electroporated with multiplexed sgRNA against *TRIM5*. Single cell clonal lines were generated by limiting dilution to generate a THP-1 *TRIM34 TRIM5* double KO clonal cell line. Cells were doubly transduced with doxycycline-inducible, HA-tagged human TRIM34 or empty vector control in tandem with doxycycline-inducible, FLAG-tagged human TRIM5α, TRIM5Δv1, TRIM5 R332P, or empty vector control. Expression levels were visualized by immunoblotting 72 h post-induction. b-d. THP-1 *TRIM34 TRIM5* double KO clonal cells co-expressing doxycycline-inducible human TRIM34 or empty vector control and human TRIM5α (grey bars), TRIM5Δv1 (red bars), TRIM5 R332P (blue bars), or empty vector control were infected with SIV_AGM-SAB_ (b), HIV-1 N74D (c), or HIV-1 (d) CA 1 day post-induction. Levels of infection were quantified 2 dpi by flow cytometry. Relative infectivity in induced cells (solid bars) is normalized to average infectivity in uninduced control cells (not shown). Data are represented as mean +/- s.d. of 3 technical replicates from 1 representative experiment. One-sided p values were calculated by Welch’s *t*-test. ns = nonsignificant, * p ≤ 0.05, ** p ≤ 0.01, *** p ≤ 0.001, **** p ≤ 0.0001.

Given that TRIM5 is required for TRIM34-mediated restriction, we next asked whether the TRIM5α v1 loop, known to be important for capsid specificity by TRIM5α, was essential for restriction by TRIM34. We found that unlike co-expression of TRIM34 with full-length TRIM5α, co-expression of TRIM34 with TRIM5α Δv1 was not sufficient for restriction of either SIV_AGM-SAB_ and HIV-1 N74D (Figure 3b-c; red bars). This suggests that the TRIM5α v1 loop is required for TRIM34-mediated restriction.

We then asked whether changing the identity of the TRIM5α v1 loop, but not deleting it entirely, could also alter viral specificity. To assess this question, we generated the mutant TRIM5α R332P, which converts the human residue at position 332 to the rhesus macaque residue and has been shown to have enhanced antiviral inhibition of HIV-1 [21,30]. We co-expressed these cells with TRIM34 and infected with SIV_AGM-SAB_, HIV-1 N74D, or HIV-1. In accordance with the literature, TRIM5α R332P was able to restrict HIV-1 while WT TRIM5α did not (Figure 3d; blue bars) [21,30]. Furthermore, restriction of HIV-1 by TRIM5α R332P was agnostic to the presence of TRIM34; that is, TRIM34 did not appear to augment restriction of HIV-1 by TRIM5α R332P (Figure 3d; blue bars). Similar to wild type TRIM34 and TRIM5α, TRIM34 co-expression with the mutant TRIM5α R332P allele maintained restriction of the N74D CA mutant virus (Figure 3c; blue bars). Interestingly, TRIM5α R332P alone did not restrict HIV-1 N74D, in contrast to WT HIV-1 (Figure 3c-d; blue bars). This suggests that TRIM34 is critical for restriction of HIV-1 N74D, even in the presence of a TRIM5α mutant that is more potent against WT HIV-1. Finally, TRIM34 co-expressed with TRIM5α R332P restricted SIV_AGM-SAB_ CA, whereas TRIM5α R332P alone did not (Figure 3b; blue bars). These findings support a role for both the TRIM5α v1 loop and for TRIM34 in viral recognition and specificity.

### TRIM34 SPRY Domain is Required for Restriction

To more directly test whether the TRIM34 SPRY domain plays a role in antiviral activity and specificity, we generated a chimeric protein that encodes the TRIM34 RING, Bbox and coiled-coil domains in frame with the TRIM5 SPRY domain (TRIM34 RBCC-TRIM5 SPRY) (Figure 4a). We hypothesized that if the TRIM34 SPRY domain was required for restriction, the chimera would lose restriction relative to full-length TRIM34. We transduced this chimera into THP-1 *TRIM34* clonal KO cells and confirmed expression by Western blot (Figure 4b). We assayed this construct for restriction in our THP-1 cells that lack TRIM34 expression but do contain endogenous, full-length TRIM5α. We found that in the presence of endogenous TRIM5α, the TRIM34 RBCC-TRIM5α SPRY chimera was not able to restrict SIV_AGM-SAB_ CA (Figure 4c). Therefore, the TRIM34 SPRY domain, in addition to the TRIM5α SPRY domain, is involved in restriction.

**Figure 4.**
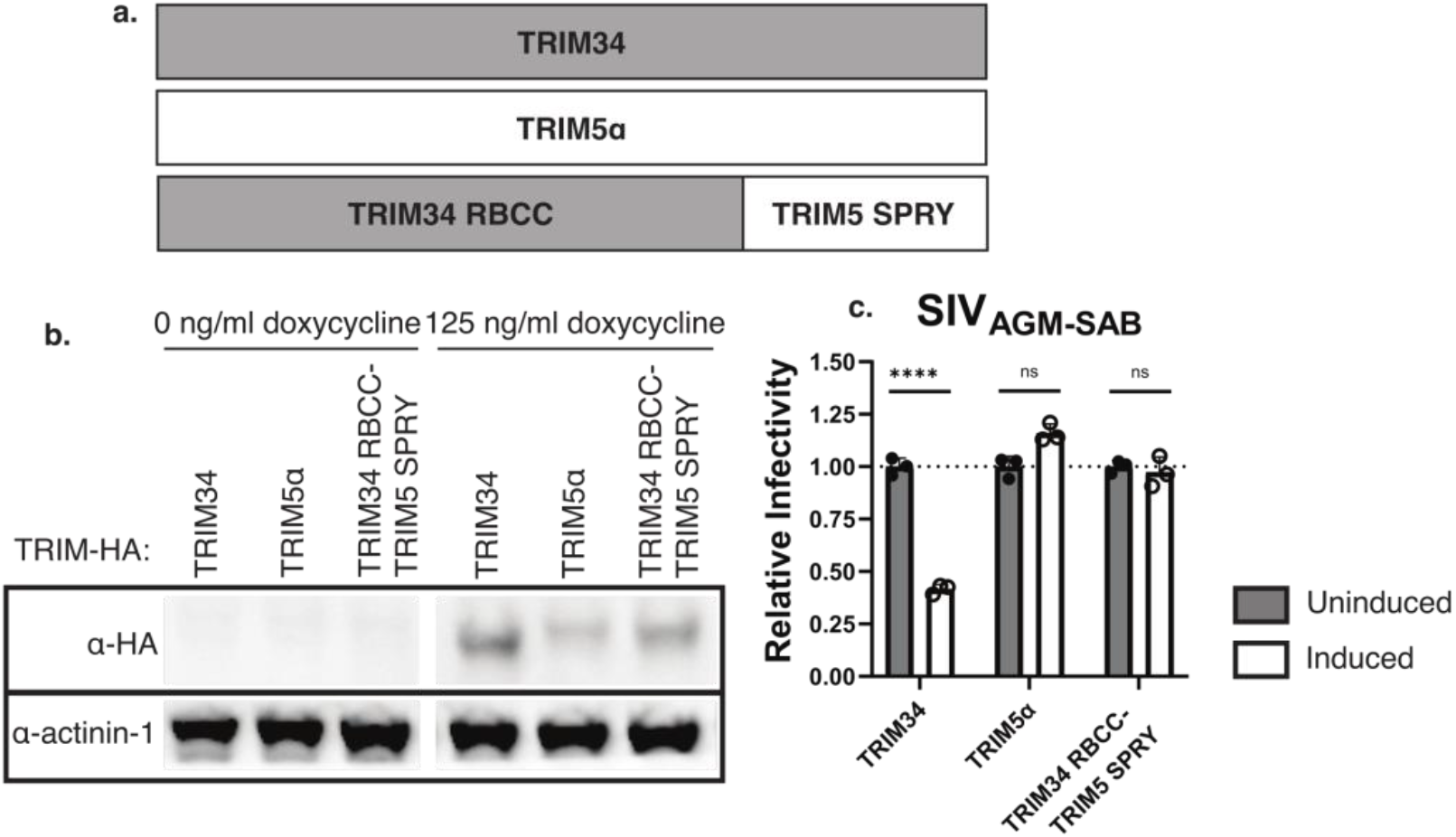
TRIM34 SPRY domain is necessary for restriction of SIV_AGM-SAB_. a. Schematic of TRIM34 RBCC-TRIM5 SPRY chimera b. THP-1 *TRIM34* clonal KO cells were generated by electroporation of multiplexed sgRNA against *TRIM34* and single cell sorting. Cells were transduced with doxycycline-inducible HA-tagged human TRIM34, TRIM5α, or TRIM34-TRIM5α chimeras. Expression levels were visualized by immunoblotting 72 h post-induction. c. THP-1 *TRIM34* clonal KO cells expressing doxycycline-inducible human TRIM34, TRIM5α, or TRIM34 RBCC-TRIM5 SPRY were infected with SIV_AGM-SAB_ CA particles 1 day post-induction. Level of infection was quantified 2 dpi by flow cytometry. Relative infectivity in induced cells (white bars) is normalized to average infectivity in uninduced control cells (grey bars). Data are represented as mean +/- s.d. from 1 representative experiment comprising 3 technical replicates. One-sided p values were calculated by Welch’s *t*-test. ns = nonsignificant, * p ≤ 0.05, ** p ≤ 0.01, *** p ≤ 0.001, **** p ≤ 0.0001.

## DISCUSSION

Human TRIM34 was recently identified as a restriction factor of the CA mutant HIV-1 N74D and certain SIVs [16]. Here we show that this antiviral phenotype is a broadly conserved function of primate *TRIM34* genes. We find that the same subset of lentiviral capsids is restricted by a diverse panel of TRIM34 orthologues, irrespective of TRIM34 species of origin. Moreover, we find that although antiviral specificity does not seem to vary with the identity of TRIM34, both the TRIM34 and TRIM5α SPRY domains are critical to restriction, suggesting that TRIM5 confers a level of specificity to the TRIM34 restriction.

Despite the lack of evidence for positive selection on TRIM34, primate TRIM34 alleles have broadly maintained the ability to restrict certain primate lentiviruses. This is surprising in light of the fact that positive selection is a marker of critical regions of direct host-viral interaction. We propose that although TRIM34 lacks this characteristic, it utilizes TRIM5α, which is a rapidly-evolving gene in primates, to restrict lentiviruses. Prior work has shown that TRIM34 and TRIM5α are capable of multimerizing *in vitro* in addition to binding capsid [9,13,14]. In this work we demonstrate that TRIM5α can functionally interact with a number of primate TRIM34 orthologues. This supports a model in which TRIM5α and TRIM34 multimerize to form a higher-order structure that enables both SPRY domains to interact with the viral capsid. Therefore, despite the lack of an overt, positively-selected site of viral interface on TRIM34, TRIM34 restricts viral infection through interaction with the rapidly-evolving TRIM5 protein. It is also possible that physical interaction between TRIM34 and TRIM5α alters the half-life of one or boTHProteins, affecting turnover rate or stability. Prior work has demonstrated that different TRIMs have different half-lives, and that alterations to the RING or Bbox domains of TRIM proteins can affect protein turnover [14,31,32]; thus, it is possible that interactions between different TRIMs might lead to changes in stability.

Another possibility is that TRIM34 could be filling the role of an effector molecule. Previously, the TRIM5α RING domain has been implicated as an E3 ubiquitin ligase that acts as a signal transducer to initiate innate immune activation subsequent to CA sensing by the TRIM5α SPRY domain [33–35]. In the case of TRIM34, it is possible that TRIM5α is responsible for recognizing CA while TRIM34’s E3 ligase domain contributes to downstream signaling, capsid degradation, or recruitment of other molecules that aid restriction. Although Lascano *et al*. found that TRIM34 does not activate AP-1, it remains possible that it is acting by another mechanism [34]. Notably, these models are not mutually exclusive and could function in tandem with TRIM34 aiding TRIM5α in more than one capacity.

Our findings suggest that the SPRY domains of both TRIM34 and TRIM5α are important for restriction. On its own, human TRIM5α is a relatively weak restriction factor. We propose that TRIM34 can assist human TRIM5α to restrict some—but not all—viral capsids. For example, TRIM5α in the presence of TRIM34 is able to restrict SIV_AGM-SAB_; conversely, even in the presence of TRIM34, TRIM5α is still unable to restrict HIV-1. One possibility is that the TRIM34 SPRY domain is contributing to viral recognition. Previous work has shown that TRIM34 is able to bind HIV-1 CA-NC complexes *in vitro*, even though it does not restrict HIV-1, raising the possibility that TRIM34 may still contribute to capsid binding [14,36]. Although the v1 loop of TRIM34 has not undergone positive selection and is only 6 amino acids long, we cannot exclude the possibility that it, or one of the other variable loops, might contribute to capsid recognition. Given the relatively high conservation of the variable loops of the primate TRIM34 alleles that we assessed (no amino acid differences in v1, v3, or v4; 1 amino acid difference in each allele in v2), this could explain the observation that all the TRIM34 alleles tested restricted the same subset of viruses. Furthermore, the observation that TRIM5α R332P—which gains restriction of HIV-1 relative to WT TRIM5α—does not restrict HIV-1 N74D alone and only restricts it in the presence of TRIM34, supports the idea that TRIM34 could be assisting in capsid recognition beyond the capacity of TRIM5α alone. Thus, as many TRIM5α alleles can act as potent restriction factors of a few very specific viral capsids, but a poor restriction factor of many others, we speculate that TRIM34 might act as a cofactor to enable TRIM5α to restrict viral capsids that TRIM5α is not able to restrict on its own.

It is intriguing that all the TRIM34 orthologues tested restricted the same subset of viruses: SIV_AGM-SAB_, SIV_AGM-TAN_, and SIV_MAC_. Notably, only viruses originating from Old World monkeys were restricted, whereas the hominid viruses, HIV-1 and SIV_CPZ_, were not restricted. It is possible that this is driven by the identity of TRIM5α: HIV-1 and SIV_CPZ_ are better evolved to evade restriction by human TRIM5α compared to the Old World monkey viruses [23]. The three restricted viruses share only 70.6% pairwise amino acid identity across their CA regions, making it difficult to identify critical residues or motifs that determine restriction. We explored the possibility that restriction might be dependent on CypA binding. However, CypA binding capacity varied across the restricted viruses. For example, SIV_AGM-TAN_ does bind CypA, while SIV_MAC_ and SIV_AGM-SAB_ do not [13,37–39]. Furthermore, while HIV-1 is well-established to bind and incorporate CypA into virions, the CypA binding capacity of HIV-1 N74D is controversial [15,40,41]. Thus, at minimum, restriction does not depend exclusively on CypA binding. Instead, restriction of these capsids by TRIM34 and TRIM5α may be dependent on other unknown features that distinguish these capsids. A number of blocks to HIV infection that are capsid-dependent have been characterized—for example, Lv2, Lv3, and Lv4—for which the responsible cellular components have either not been identified or have only been partly identified [42–46]. It is possible that TRIM34 could contribute to one or more of these blocks.

We find that TRIM34 alleles from a broad spectrum of primates, when paired with TRIM5α, are able to restrict capsids that neither TRIM is able to restrict on its own. Despite lacking signs of positive selection characteristic of many restriction factors, TRIM34—acting in tandem with TRIM5α—can act as a barrier to cross-species transmission events. It is possible that these proteins have co-evolved such that TRIM34 can enhance or modify TRIM5α’s antiviral potential. Indeed, not only do host immune proteins evolve in the context of the pathogens they counteract, but they also evolve in the context of other, complementary host proteins. Thus, our data suggest that restriction factors evolve not only in isolation in response to evolutionary pressures exerted by viral pathogens but may also co-evolve with each other resulting in more powerful antiviral activity than either could achieve on its own.

## ACKNOWLEDGEMENTS

We thank Jeannette Tenthorey and Vanessa Montoya for critical feedback of this manuscript. We thank Melissa Kane for sharing the doxycycline-inducible vector system, Jeannette Tenthorey for the TRIM5 mutant constructs, and Theodora Hatziioannou for the chimeric capsid constructs. This work was supported by the Viral Pathogenesis and Evolution Training Grant (NIH T32 AI083203) awarded to J.T., DP1 DA051110 to M.E., and NIH/NIAID R01 AI147877 and U54 AI170856, to M.O. This research was supported by the Genomics, Bioinformatics, and Flow Cytometry Shared Resources, RRID:SCR_022606, of the Fred Hutchinson/University of Washington Cancer Consortium (P30 CA015704).

## AUTHOR CONTRIBUTIONS

Conceptualization, methodology, writing, and funding acquisition were performed by J.T., M.E., and M.O. Investigation was conducted by J.T., A.K., and A.L.F. Validation, formal analysis, and visualization were performed by J.T. and A.K. Supervision was provided by M.E. and M.O.

## MATERIALS AND METHODS

### Cell culture

All cells were cultured at 37°C and 5% CO_2_. THP-1 monocytic cells (American Type Culture Collection, Manassas, VA, #TIB-202) were cultured in Roswell Park Memorial Institute 1640 Media (RPMI 1640) (Gibco, Grand Island, NY, #118875-093) supplemented with 10% v/v fetal bovine serum (FBS) (GE Cytiva, Marlborough, MA #SH30541.03), 100 U/mL penicillin-streptomycin (Gibco #15140-122), 10mM HEPES (Gibco #15630-080), 1mM sodium pyruvate (Gibco #11360-070), 2 g/L D-Glucose (Gibco #A24940-01), and 1X GlutaMAX supplement (Gibco #35050-061). HEK 293T/17 cells (American Type Culture Collection #CRL-11268) and HeLa cells (American Type Culture Collection #CCL-2) were cultured in Dulbecco’s Modified Eagle Medium (DMEM) (Gibco #11965-092) supplemented with 10% fetal bovine serum, and 100 U/mL penicillin-streptomycin.

### Cloning, plasmids, and virus production

All transfections were performed on HEK 293T/17 cells in the presence of serum-free DMEM and Trans-IT transfection reagent (Mirus Bio, Madison, WI, #MIR 2305). All transductions were performed by spinoculation at 1100 x *g* for 30 min at 30°C.

Fluorescent reporter viruses were generated by transfection of a three-plasmid system: pMD2.G (Addgene, Watertown, MA, #12259, a gift from Didier Trono) for expression of VSV-G envelope; a variable plasmid for expression of a chimeric gag/pol containing an NL4-3 backbone and HIV-1 [22], SIV_CPZ_, SIV_AGM-SAB_, or SIV_MAC_ CA (gifts from Theodora Hatziioannou) [23]; and either pALPS-eGFP (Addgene #101323, a gift from Jeremy Luban) [47] or pHIV-zsGreen (Addgene #18121, a gift from Bryan Welm and Zena Werb) [48]. SIV_CPZ_, SIV_AGM-SAB_, or SIV_MAC_ chimeric virus particles were made wiTHPALPS-eGFP, and HIV-1 virus particles were made wiTHPHIV-zsGreen. The SIV_MAC_ CA construct contained the mutation A77V. Luciferase reporter viruses were generated by transfection of a two-plasmid system: SIVagmTAN E-R-luc [24] and L-VSV-G (VSV glycoprotein expression) [49]. Particles were harvested 2 days post-transfection, syringe-filtered through 0.22uM PES membranes, and frozen at −80°C. N-MLV reporter viruses were generated by transfection of a three-plasmid system: pCIG3-N for expression N-MLV gag/pol (Addgene #132941, a gift from Jeremy Luban) [50], pQCXIP-eGFP (a gift from Jeannette Tenthorey) [21] for expression of a fluorescent reporter, and pMD2.G for expression of VSV-G envelope. NB-MLV reporter viruses were generated by transfection of a four-plasmid system: JK3 for expression of NB-MLV gag/pol [49], L-VSV-G for expression of VSV-G envelope [49], CMV-tat for transactivation [49], and pQCXIP-eGFP for expression of a fluorescent reporter.

pLentiCRISPR-v2 (Addgene #52961, a gift from Feng Zhang) constructs were generated by BsmBI (New England BioLabs, Ipswich, MA, #R0580) restriction cloning of TRIM5 guides into pLentiCRISPR-v2, a lentiviral vector encoding for Cas9 [51]. pLentiCRISPR-v2 containing guides were transfected wiTHPSPAX2 (Addgene #12260, a gift from Didier Trono) and pMD2.G to generate lentiviral particles. Particles were harvested 2 days post-transfection, syringe-filtered through 0.22 μM PES membranes, and frozen at −80°C.

HA-tagged TRIM34 and 3xFLAG-tagged TRIM5α codon-optimized expression constructs (Supplemental data) were synthesized by Twist Bioscience (South San Francisco, CA). TRIM5α mutant constructs (TRIM5α v1 loop and TRIM5α R332P) were a gift from Jeannette Tenthorey [21]. TRIM constructs were cloned into pLKO (a gift from Melissa Kane) [52] by SfiI (New England BioLabs #R0123S) restriction enzyme digest. pLKO-TRIM constructs were transfected along wiTHPMD2.G and pSPAX2 to generate particles. Particles were harvested at 2 and 3 days post-transfection, cell pellets spun down at 300 x *g*, and supernatants frozen at −80°C.

HA-tagged TRIM34ΔSPRY constructs were cloned in pQCXIP (TaKaRa Bio, San Jose, CA, #631516) by SbfI (New England BioLabs #R3642S) and NotI (New England Biolabs #R3189S) restriction digest. pQCXIP-TRIM34ΔSPRY constructs were transfected along wiTHPJK3 (MLV gag/pol), L-VSV-G (VSV-G envelope), and CMV-tat to generate particles. Particles were harvested 2 days post-transfection.

### CRISPR Knockouts

Clonal knockout lines in THP-1 cells were generated by electroporation of multiplexed small guide RNA (sgRNA) from Gene Knockout Kit v2 (Synthego, Redwood City, CA) against *TRIM34* (guide sequences = CTTGCTTAACGTACAAG, CCACAGTCTAGACTCAA, GCAGTGACCAGCATGGG) or *TRIM5* (guide sequences = GGUAACUGAUCCGGCACACA, ACUUCUUGUGGUUUGCAGUG, CCUGGUUAAUGUAAAGGAGG). Single cell clonal lines were generated by single cell sorting (*TRIM34*) or limiting dilution (*TRIM5*). Knockout efficiency was validated by Interference of CRISPR Edits (ICE) analysis (Synthego) [53].

Pooled knockout lines were generated by transduction of THP-1 cells with lentiviral preps containing guides delivered by pLentiCRISPR-v2. *TRIM5* guide sequences = TCACCACACGTTCCTCACAG and GTTGATCATTGTGCACGCCA. NTC guide sequences = GGGCCCGCATAGGATATCGC and TAGACAACCGCGGAGAATGC. Cells were spinoculated in the presence of 20 μg/mL DEAE-Dextran (Pharmacia Fine Chemicals, Uppsala, Sweden, #17-0350-01) and then selected in 10 μg/mL blasticidin S HCl (Gibco #A11139-03). Knockout efficiency was validated by ICE analysis [53].

### Inducible Overexpression

Doxycycline-inducible expression of TRIM34 and TRIM5α was achieved by transduction of lentiviral preps containing pLKO TRIM constructs in THP-1 cells. Cells were spinoculated in the presence of 5 μg/mL polybrene (EMD Millipore, Burlington, MA, #TR-1003-G) and then selected in 0.5 μg/mL puromycin (TRIM34) (Sigma-Aldrich, St. Louis, MO, #P8833-25MG) or 10 μg/mL blasticidin (TRIM5α). Protein expression was induced by the addition of 125 ng/mL doxycycline hyclate (Sigma-Aldrich #D9891-5G) to cells cultured in complete RPMI 1640 containing tetracycline-approved FBS (Sigma-Aldrich #F0926-50ML).

Stable expression of TRIM34ΔSPRY was achieved by transduction of lentiviral preps containing pQCXIP constructs expressing TRIMΔSPRY constructs. HeLa cells were spinoculated in the presence of 20 μg/mL DEAE-Dextran. Selection was performed in the presence of 1 μg/mL puromycin.

### Restriction assays

1 day after induction of TRIM expression, THP-1 cells were infected with chimeric CA virus particles expressing a fluorescent reporter. Cells were spinoculated in the presence of 20 μg/mL DEAE-Dextran. 2 dpi, relative infectivity was quantified by flow cytometry using a FACSCelesta Analyzer (BD Biosciences, San Jose, CA) or Bright-Glo luciferase assay reagent (Promega, Madison, WI #E2620) using a LUMIstar Omega luminometer (BMG Labtech, Ortenberg, Germany).

### N-MLV Restriction assay

HeLa cells that had been transduced to stably express TRIM34ΔSPRY constructs were infected with N-MLV particles expressing an eGFP reporter by spinoculation in the presence of 20 μg/mL DEAE-Dextran. Infectivity was assessed 2 dpi by flow cytometry using an LSRFortessa Cell Analyzer (BD Biosciences).

### Western blotting

3 days after induction of TRIM expression, THP-1 cells were harvested; 1 day prior to infection, HeLa cells were harvested. Cells were washed once wiTHPBS. Cells were then lysed on ice for 30 min in 20 mM HEPES (Fisher Scientific, Fair Lawn, NJ, #BP310-1) containing 8 M urea (Sigma-Aldrich #U-6504), 50 mM DL-Dithiothreitol (DTT) (Gold Biotechnology, St. Louis, MO, #DTT10), 0.1% w/v SDS (Fisher Scientific #BP166-500), 1.5 mM MgCl2 (Sigma-Aldrich #208337-100G), 0.5 mM CaCl2 (Sigma-Aldrich #C-3306), 50 μg/mL DNAse I (Roche Diagnostics, Mannheim, Germany, #10104159001), and 1X EDTA-free protease inhibitor cocktail (Roche Diagnostics #11836170001). 6X SDS-PAGE sample loading buffer (G Biosciences, St. Louis, MO #785-701) containing 5% v/v 2-mercaptoethanol (Sigma-Aldrich #M3148-100ML) was added to lysates. Lysates were acid treated with 50 mM HCl (Sigma-Aldrich #320331-300ML) and then boiled at 95°C for 7 min. After cooling, lysates were neutralized with 50 mM NaOH (Sigma-Aldrich #72068-100ML). Products were resolved by SDS-PAGE on NuPage Bis-Tris 4-12% acrylamide gels (Invitrogen, Carlsbad, CA, #NP0335BOX) and transferred onto nitrocellulose membranes (Bio-Rad Laboratories, Hercules, CA, #1620115). Membranes were blocked with 2% w/v milk (Research Products International, Mt. Prospect, IL, #M17200-500.0) and 2% w/v bovine serum albumin (Sigma-Aldrich #A7906-100G) in tris-buffered saline (Fisher Scientific #BP152-5) containing 1% v/v Tween-20 (Fisher Scientific #BP152-1) (TBS-T). Primary antibodies were incubated overnight at 4°C: rabbit anti-HA high affinity at 1:500 (Roche Diagnostics #ROAHAHA), mouse anti-FLAG M2 at 1:500 (Sigma-Aldrich #F1804), and rabbit anti-alpha actinin-1 at 1:1000 (Bio-Rad Laboratories #VPA00889). Membranes were washed with TBS-T and then incubated with HRP-conjugated secondary antibodies for 1 h at RT: goat anti-rat IgG at 1:2000 (Abcam, Waltham, MA #ab97057), sheep anti-mouse IgG at 1:2000 (GE Cytiva #NA931), and donkey anti-rabbit IgG at 1:2000 (GE Cytiva #NA934V). Membranes were washed again and then incubated with SuperSignal West Pico substrate (HA and alpha actinin-1 blots) (Thermo Fisher Scientific, Waltham, MA, #34580) or Supersignal West Femto substrate (FLAG blots) (Thermo Fisher Scientific #34095) for 3 min. Membranes were imaged with a ChemiDoc MP imaging system (Bio-Rad Laboratories).

### Statistical methods

For comparisons of 2 independent variables, one-tailed p values were computed by Welch’s *t*-test. For comparisons of 3 or more independent variables, p values were computed by Brown-Forsythe and Welch’s 1-way ANOVA with Dunnett’s T3 test for multiple comparisons. All calculations and data visualizations were performed using GraphPad Prism version 9.3 (GraphPad Software, San Diego, CA).

## ADDITIONAL FILES

Additional File 1 (Additional File 1.txt). TRIM34 and TRIM5a sequences used in this study, codon-optimized for expression in human cells where applicable.

